# Complex human gut microbiome cultured in anaerobic human intestine chips

**DOI:** 10.1101/421404

**Authors:** Sasan Jalili-Firoozinezhad, Francesca S. Gazzaniga, Elizabeth L. Calamari, Diogo M. Camacho, Cicely W. Fadel, Bret Nestor, Michael J. Cronce, Alessio Tovaglieri, Oren Levy, Katherine E. Gregory, David T. Breault, Joaquim M. S. Cabral, Dennis L. Kasper, Richard Novak, Donald E. Ingber

**Author notes:** These authors contributed equally to this work (Sasan Jalili-Firoozinezhad, Francesca S. Gazzaniga, Elizabeth L. Calamari). Corresponding Author: Donald E. Ingber.

## Abstract

The diverse bacterial populations that comprise the commensal microbiota of the human intestine play a central role in health and disease, yet no method is available to sustain these complex microbial communities in direct contact with living human intestinal cells and their overlying mucus layer *in vitro*. Here we describe a human Organ-on-a-Chip (Organ Chip) microfluidic platform that permits control and real-time assessment of physiologically-relevant oxygen gradients, and which enables co-culture of living human intestinal epithelium with stable communities of aerobic and anaerobic human gut microbiota. When compared to aerobic co-culture conditions, establishment of a transluminal hypoxia gradient sustained higher microbial diversity with over 200 unique operational taxonomic units (OTUs) from 11 different genera, and an abundance of obligate anaerobic bacteria with ratios of *Firmicutes* and *Bacteroidetes* similar to those observed in human feces, in addition to increasing intestinal barrier function. The ability to culture human intestinal epithelium overlaid by complex human gut microbial communities within microfluidic Intestine Chips may enable investigations of host-microbiome interactions that were not possible previously, and serve as a discovery tool for development of new microbiome-related therapeutics, probiotics, and nutraceuticals.

---

One of the major recent paradigm shifts in medicine relates to the recognition of the central role that the microbiome, composed of host-specific communities of commensal microbes, plays in human health and disease^1^. While human microbiota colonize mucosal surfaces of various tissues, the gastrointestinal tract supports the greatest mass and diversity of microorganisms^2^. Aerobic and anaerobic commensal gut microbiota are essential for maintenance of normal nutrient absorption, drug metabolism, and immune responses, as well as for protection against infectious pathogens^3^. Conversely, changes or imbalances in the microbial community within the intestine can contribute to development of a broad range of pathological disorders within and beyond the gastrointestinal system, including inflammatory bowel disease, colorectal cancer, radiation enteropathy, diabetes, hepatic steatosis, obesity, and rheumatoid arthritis^4,5^. Thus, the establishment and preservation of balanced host-intestinal microbiome interactions are key requirements for maintaining gut homeostasis and human health.

Analysis of gut-microbiome crosstalk has almost exclusively relied on genomic or metagenomic analysis of samples collected *in vivo* because no method exists to establish stable complex communities of gut commensal microbes in direct contact with intestinal epithelium and their overlying mucus layer *in vitro*^6,7^. While animal models have been used to analyze host-microbiome interactions and their contributions to pathophysiology^8–10^ there are no *in vitro* systems available to verify these interactions in human cells when cultured in direct contact with complex human microbiome. Thus, there is a great need for experimental models that can sustain complex populations of human aerobic and anaerobic microbiota in contact with living human tissues to analyze dynamic and physiologically relevant human host-microbiome interactions.

Existing *in vitro* models, such as Transwell inserts, have been used to study human host-microbe interactions; however, these studies can only be carried out over a period of hours before bacterial overgrowth leads to cell injury and death^11–13^. Organoid cultures, have shown great promise for studying host-microbiome interactions, but they also cannot be co-cultured with living microbes for more than ∼24 hours, they do not provide a vascular interface, nor can they sustain luminal oxygen levels below 0.5% as is required for co-culture of certain obligate anaerobes^14,15^. Specialized bioreactor models, such as the mucosal-simulator of the human intestinal microbial ecosystem (M-SHIME), have been developed to sustain growth of luminal and mucosal gut microbes *in vitro*, but they do not include living human intestinal epithelium, and hence, they cannot be used to study host-microbiome interactions^16^. We previously described a microfluidic Organ Chip device lined by human Caco2 intestinal epithelial cells cultured under dynamic fluid flow and peristalsis-like mechanical deformations, which enabled establishment of stable co-cultures of a human villus intestinal epithelium in direct contact with up to 8 different strains of human commensal gut microbes for weeks *in vitro* under aerobic conditions^17–19^, but the human gut microbiome contains hundreds of different types of bacteria, many of which are obligate anaerobes that will not grow in this environment. A human-microbiota interaction (HMI) module also has been developed that permits analysis of aerobic and anaerobic microbes, including complex living microbiome derived from a SHIME reactor, when co-cultured with human Caco2 intestinal epithelial cells under an oxygen gradient; however, the microbes were separated from the human cells by a nanoporous membrane with an artificial mucus layer, and even under these conditions, the co-cultures were only maintained for 48 hours^17^. Thus, no existing *in vitro* model enables analysis of direct interactions between commensal gut bacteria and human intestinal epithelium through its overlying mucus layer when cultured for multiple days *in vitro*, which is crucial for analyzing gut health and disease^20–22^.

In this study, we therefore set out to develop an experimental system that can support dynamic interactions between living, mucus-producing, human intestinal epithelium and a directly apposed complex community of living human aerobic and anaerobic commensal gut microbes with a population diversity similar to that observed in living human intestine. To meet this challenge, we modified the human Caco2 Intestine Chip by integrating microscale oxygen sensors into the devices for *in situ* oxygen measurements, and placing the chips within an engineered anaerobic chamber to establish a physiologically relevant oxygen gradient across a vascular endothelium and intestinal epithelium that are cultured in parallel channels separated by a porous membrane within the device. To insure a stable source of complex human intestinal gut-microbiota, we used complex microbiota originally derived from healthy human stool specimens, which have been maintained stably in gnotobiotic mice for multiple years, and closely resemble the relative abundance of major bacterial phyla patterns in their respective inoculum^23,24^. We also applied the same method to co-culture fresh gut microbiome isolated from human infant stool samples in a primary human Intestine Chip lined by cells isolated from normal human ileum. Here we describe how establishing a hypoxia gradient across the engineered tissue-tissue (endothelium-epithelium) interface of the Intestine Chip allows us to stably co-culture complex communities of anaerobic and aerobic human commensal gut bacteria in the same channel as human villus intestinal epithelium while simultaneously monitoring oxygen levels for at least 5 days *in vitro*.

## RESULTS

### Establishing an oxygen gradient across the lumen of the Intestine Chip

To recapitulate a physiologically relevant intestinal oxygen gradient profile inside Intestine Chips (**Fig. 1a**), we fabricated an oxygen-sensing, dual channel, human Organ Chip composed of optically clear and flexible poly(dimethyl siloxane) (PDMS) polymer (**Fig. 1b; Supplementary Fig. S1a**), as well as an anaerobic chamber. For real-time, non-invasive, monitoring of oxygen tension, six sensor spots containing oxygen-quenched fluorescent particles were embedded in the top and bottom portions of the chip beneath the central microchannels (**Fig. 1b; Supplementary Fig. S1b**). Changes in the fluorescent intensities of these sensors in response to oxygen tension (**Supplementary Fig. S1b**), were captured by a VisiSens camera, and translated into oxygen concentrations by comparison with a standard Oxy-4 probe system (**Supplementary Fig. S1c**). As both the chips and sensors are composed of highly gas-permeable PDMS, the sensors respond rapidly (< 30 sec) to changes in oxygen concentrations (**Fig. 1c**).

**Figure 1.**
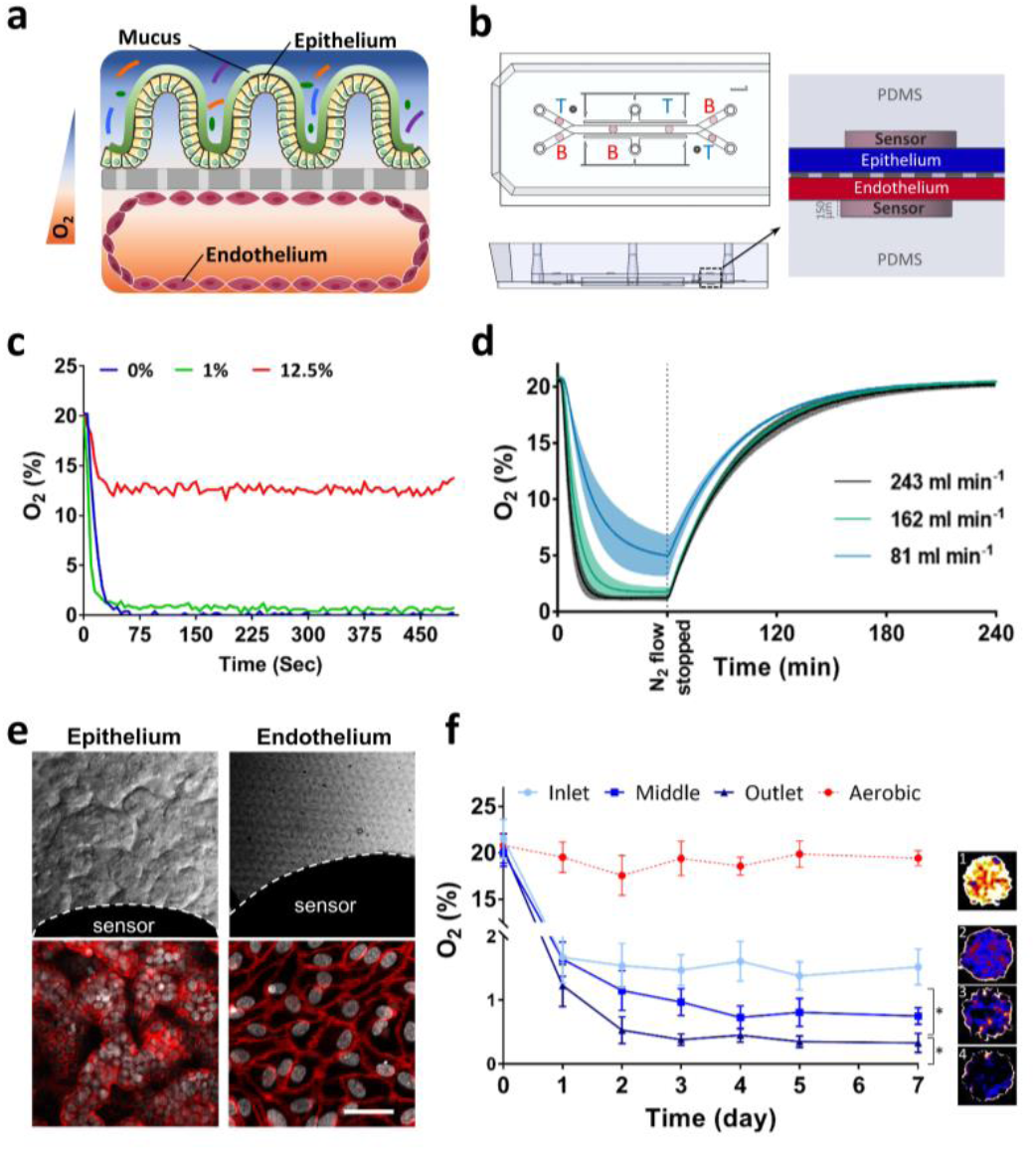
Oxygen sensitive human Intestine Chip microfluidic culture device. (**a**) Schematic showing the position of the human intestinal epithelium overlaid with its own mucus layer and complex gut microbiota on top, with vascular endothelium on bottom side of the same ECM-coated porous membrane, within a 2-channel microfluidic Organ Chip device in presence of oxygen gradients. Orange and blue colors indicated high and low levels of oxygen concentration, respectively. (**b**) Schematic representation of the Intestine Chip with 6 oxygen quenched fluorescent particles embed in inlet, middle and outlet of top and bottom channels. (T, top channel; B, bottom channel). (**c**) Sensitivity analysis of oxygen spots located in the Intestine Chip in response to defined, standard oxygen concentrations. (**d**) Anaerobic chamber validation at various N2 inflow pressures; N2 was introduced into the chamber at 81 mL min^-1^, 162 mL min^-1^, or 243 mL min^-1^ for 1 h before gas flow was stopped and the chamber was allowed to recover (n=3, shaded regions indicate standard deviation). (**e**) Microscopic views showing the villus morphology of the human Caco-2 intestinal epithelium (bar, 100 μm) and vascular endothelium (bottom left; bar, 100 μm) cultured for 6 days in the Intestine Chip under anaerobic conditions, when viewed from above by DIC and phase contrast imaging, respectively, or by immunofluorescence staining for the tight junction protein, ZO-1 (red, top right; bar, 100 μm) and endothelial cell junction-associated protein, VE-cadherin (red, bottom right; bar, 20 μm). Gray indicates DAPI-stained nuclei; white dashed lines indicate the border of the oxygen sensor spot). (**f**) Oxygen concentration profiles within aerobically- and anaerobically-cultured Intestine Chips (^*:^ P<0.05). Representative pseudocolor insets indicate average oxygen concentration in aerobic chip (1), and inlet (2), middle (3) and outlet (4) of the anaerobically-cultured epithelium channel, at day 7 of culture.

To simultaneously provide adequate oxygen for maintaining human cells and an anaerobic microenvironment suitable for culturing complex human microbiota while establishing a functional host-microbiome interface, we flushed the custom anaerobic chamber continually (**Supplementary Fig. S2a,b**) with humidified 5% CO2 in nitrogen gas. This setup enables us to maintain low oxygen levels within the lumen of the upper chamber (**Fig. 1d**), while the epithelium is sustained via diffusion of oxygen through the permeable PDMS membrane from the well-oxygenated medium flowing through the lower endothelium-lined vascular channel from external oxygenated medium reservoirs (**Supplementary Fig. S2a,b**). Using this method, anaerobic conditions (<0.5%) can be generated within less than 30 min at 243 ml min^-1^ of nitrogen flow into the anaerobic chamber (**Fig. 1d**). The chamber also can sustain these low oxygen levels (<5.0%) for about 15 min after it is disconnected from the nitrogen source (**Fig. 1d**). This allows the chamber to be temporarily moved from the incubator for imaging or into a bacterial glove box (*e.g.* to replenish culture medium or add microbiota) without significantly disturbing the low oxygen environment.

When human Caco-2 intestinal epithelial cells are cultured for 5 to 7 days under aerobic conditions and dynamic flow, they undergo villus differentiation and express multiple features of the ileum portion of the human small intestine, including secretion of a mucus layer overlying the apical surface of the epithelium and establishment of barrier function^25,20,26^. Endothelial cells can also be co-cultured on the bottom of the central porous membrane in the lower channel of the same device under aerobic conditions, where they form a hollow vascular lumen lined by cells joined by VE cadherin-containing cell-cell junctions under aerobic conditions^26^. The co-culture of endothelium has been shown to enhance barrier function and mucus production (*e.g.*, expression of MUC2 and MUC5AC), as well as influence villi development and cytokine production by intestinal Caco2 epithelium under these conditions^26,27^.

When we cultured Intestine Chips lined by Caco2 intestinal epithelial cells and human intestinal microvascular endothelial cells (HIMECs) under a hypoxia gradient using our chamber, differential interference contrast (DIC) and immunofluorescence microscopic analysis confirmed that the cells again formed a villus intestinal epithelium containing polarized cells joined by ZO-1-containing tight junctions (**Fig. 1e, top; Supplementary Fig. S3a-d**) that was underlaid by a confluent HIMEC monolayer with cells linked by VE-cadherin-containing tight junctions, even under these anaerobic culture conditions (**Fig. 1e, bottom**). Both cell types also remained viable under these conditions, as measured by quantifying release of the intracellular enzyme lactate dehydrogenase (LDH), which remained relatively unchanged compared to the aerobic control during one week of anaerobic culture (**Supplementary Fig. S4a**). Quantification of the apparent permeability (*P*_*app*_) of the intestinal epithelial barrier similarly revealed no changes in the paracellular barrier function, and these human Intestine Chips displayed *P*_*app*_ values of about 1 x 10^-7^ cm s^-1^ after 7 days (**Supplementary Fig. S4b**), which are similar to those previously reported^26^. Importantly, we also confirmed that both the human intestinal epithelium and endothelium experienced oxygen gradients by analyzing the expression of hypoxia-inducible factor 1α (HIF-1α). HIF-1α is a key mediator of oxygen hemostasis and intestinal epithelial cell adaptation to oxygen deprivation,^28^ which is stabilized in a graded fashion in response to decreasing oxygen concentrations^29^. HIF-1α levels were significantly higher (∼3-fold; *p* < 0.01) in the lumen of the anaerobically-cultured epithelium than in the adjacent oxygenated endothelium-lined channel (**Supplementary Fig. S5a,b**), which is where our sensors indicated maintenance of a hypoxic environment for up to 7 days in culture (**Fig. 1f** and **Supplementary Fig. S4c**).

### Co-culture of human intestinal epithelium with an obligate anaerobe on-chip

We next explored whether the hypoxic environment can support co-culture of the intestinal epithelium with the obligate anaerobe, *Bacteroides fragilis* (*B. fragilis*; strain NCTC 9343), which is a human commensal symbiotic bacterium that cannot grow under aerobic (> 0.5% oxygen) conditions^30,31^. During the co-culture procedure which began after the epithelium had been cultured and differentiated on-chip for 7 days (**Supplementary Fig. S6a**), *B. fragilis* bacteria (2.5 x 10^5^ CFU; fluorescently labeled with HADA^32^; **Supplementary Fig. S6b**) were introduced into the lumen of the intestinal epithelium-lined upper channel (**Supplementary Fig. S6c**) and subsequently cultured under either aerobic or anaerobic conditions, while being flushed daily to remove both luminal and tissue-associated microbes and carry out CFU counts by plating.

Continuous monitoring of oxygen concentrations from inoculation to day 3 of co-culture revealed that our anaerobic chip setup maintained a low oxygen environment that decreased from ∼ 1% oxygen levels to 0.3% in the presence of *B. fragilis* (**Fig. 2a**). Yet, the intestinal epithelium maintained its ZO-1-containing tight junctions and apical brush border polarity when co-cultured in direct contact with *B. fragilis* under these highly anaerobic conditions (**Fig. 2b**). Interestingly, the presence of this obligate anaerobe enhanced barrier function (reduced P_app_ by 1.8-fold compared to aerobic conditions; *p* < 0.05) after 3 days in anaerobic culture (**Fig. 2c**) and this barrier was maintained for up to at least 8 days in culture (**Supplementary Fig. S6d**). As expected, the *B. fragilis* bacteria continued to grow in the anaerobic chips over 3 days (*p* < 0.001), whereas they started to die off and appeared at significantly lower levels under aerobic culture conditions (**Fig. 2d**). These data confirm that our chips that experience a hypoxia gradient support the growth of an obligate anaerobic bacterial species in the same channel as living human intestinal epithelial cells, whereas these bacteria would have otherwise died in a conventional aerobic culture system.

**Figure 2.**
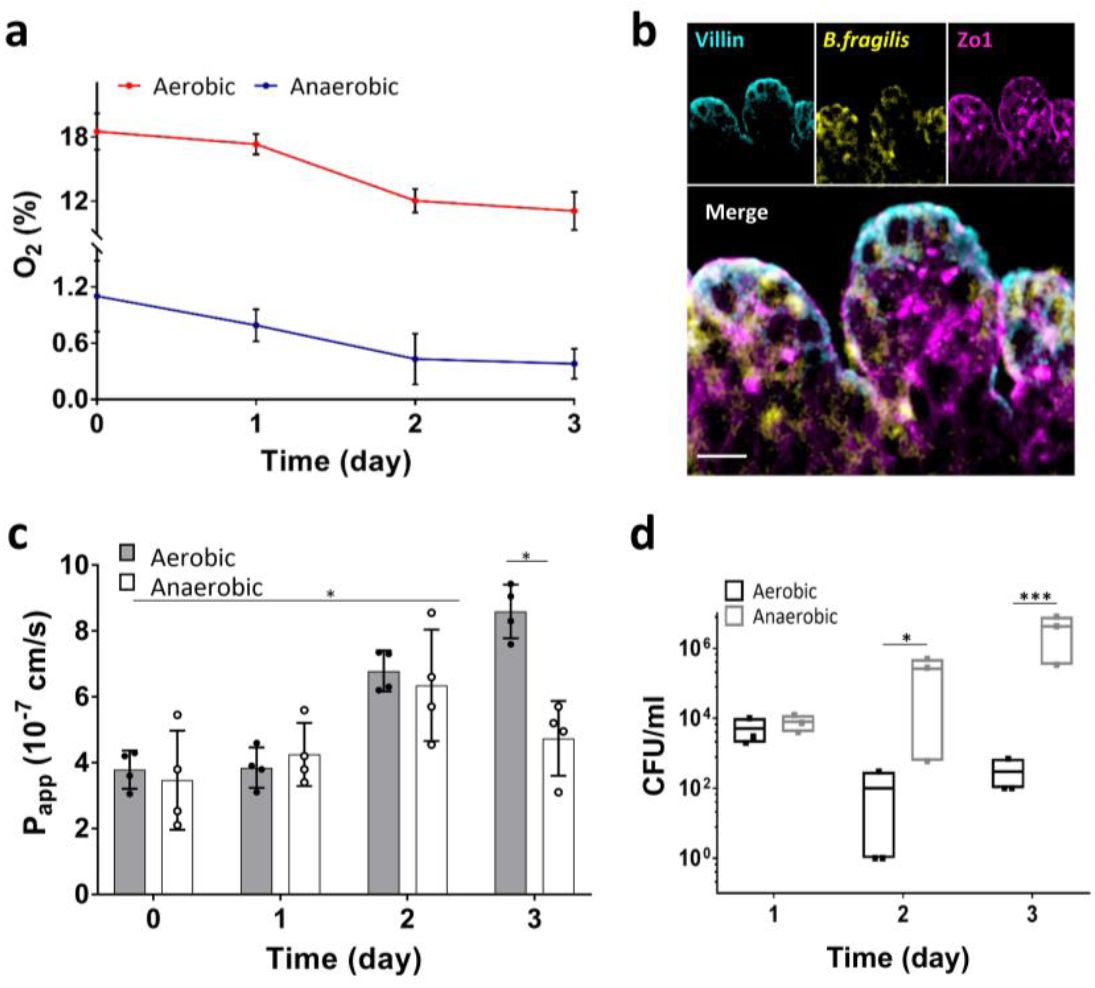
Co-culture of human intestinal epithelium and obligate anaerobe, *Bacteroides fragilis*, on-chip. (**a**) Oxygen concentration profiles in aerobic and anaerobic Intestine Chips co-cultured with *Bacteroides fragilis*. (**b**) Representative vertical cross-sectional, confocal micrographic views through the intestinal epithelium-microbiome interface within the Intestine Chip cultured under anaerobic conditions, when immunostained for villin (cyan), ZO-1 (magenta) and nuclei with DAPI (blue) (bar, 50 μm). *B. fragilis* was HADA (yellow) labeled. (**c**) Changes in apparent paracellular permeability (*P*_app_) measured by quantitating cascade blue transport across the tissue-tissue interface within the Intestine Chip microdevices co-cultured with *Bacteroides fragilis* under aerobic and anaerobic conditions (n=4; ^*:^ P < 0.05). (**d**) CFU counts/mL of *Bacteroides fragilis* co-cultured in the Intestine Chip under aerobic and anaerobic conditions (n=3; ^*:^ P < 0.05, ^***:^ P < 0.001).

### Sustaining a complex human intestinal microbiome in vitro

To optimize growth of a complex microbiome in our culture system, we searched for a source of complex human gut microbes that remains stable over time. Therefore, we inoculated the anaerobic Intestine Chips with a sample of complex gut microbiota originally isolated from human feces, which has been stably maintained in gnotobiotic mice in isolators for over 30 generations^23,24^ and that maintains a composition closely resembling the original human stool inoculum at the genera and species levels^23^. To identify a medium composition that would promote the growth of a complex set of commensal bacteria, we first inoculated the Hmb microbiota stock into 13 different types of culture medium in standard culture tubes, placed the cultures in an anaerobic chamber at 37°C, and then carried out 16S rRNA sequencing after 3 days of culture (**Supplementary Fig. S7a**). Samples of these 13 types of medium were also added to cultured human Caco2 intestinal epithelial cells to test for toxicity (**Supplementary Fig. S7b**). The medium that promoted the most diverse set of viable microbes without injuring the epithelium contained DMEM, 20% FBS, 1% glutamine, 1 mg.ml^-1^ pectin, 1 mg.ml^-1^ mucin, 5 μg.ml^-1^ Hemin and 0.5 μg.ml^-1^ Vitamin K1. The microbiota stock was introduced into this medium (0.1 mg.ml^-1^) and perfused through the upper Caco2 epithelium-lined channel of the Intestine Chip while oxygenated endothelial culture medium was flowed through the lower channel. The epithelial channel of the chips was flushed daily with a short (2 min) fluid pulse at higher flow rate (50 μl.min^-1^) to remove adherent and luminal bacteria, and 16S rRNA sequencing was carried out using samples from the effluent to assess bacterial diversity in each condition over 3 days of culture (n = 4 for each condition).

After data processing, we identified a total of 938 OTUs among all samples, which corresponded to approximately 200 unique OTUs shared between samples of each chip after filtering and removing singletons, which is similar to the scale of OTUs previously observed in human ileal aspirates (280 OTUs)^33^. Analysis of the alpha diversity between the two conditions showed that the species diversity in anaerobic chips were statistically different (PERMANOVA, *p* < 0.001) from aerobic chips (**Fig. 3a**). Although the observed diversity and Shannon Index are lower than what is observed in human stool samples (**Supplementary Fig. S8a,b**), we observed an increase in richness compared to our starting inoculum over the course of the 3 days of experiment (**Supplementary Fig. S9a,b**). We identified 11 well characterized genera including *Eubacterium, Oscillospira, Blautia, Sutterella, Biophila, Akkermansia, Ruminococcus, Bacteroides, Parabacteroides, Enterococcus and Citrobacter* (**Fig. 3b**), with an additional 8 OTUs of unknown genera from *Firmicutes* (5 OTUs) and *Proteobacteria* (3 OTUs) phyla, that were present in our chips (phylum level analysis is shown in **Supplementary Fig. S10**). Interestingly, co-culturing diverse microbiota under anaerobic conditions for 3 days in direct contact with the human intestinal epithelium did not compromise intestinal barrier integrity, and, instead, it led to an increase in barrier function by almost 2-fold (*i.e.*, decrease in P_app_ from 3.1 x 10^-7^ to 1.6 x 10^-7^ cm s^-1^ in aerobic versus anaerobic chips, respectively; *p* < 0.05) (**Fig. 3c**). In contrast, epithelial barrier function decreased (*p* < 0.001) after day 3 of co-culture under aerobic conditions when co-cultured with the same complex gut microbiome (**Fig. 3c**). Furthermore, to explore the durability of these cultures, we then carried out an additional experiment in which we extended aerobic and anaerobic chips for 5 days with and without the same Hmb stock. Sterile chips maintained barrier function over the five days of culture under both anaerobic and aerobic conditions (**Supplementary Fig. S11a,b**). Again, we observed that while barrier function decreased in aerobic chips with Hmb at day 3 (**Supplementary Fig. S11a**), no compromise of barrier function was seen in anaerobic chips with Hmb even over the 5 days of culture (**Supplementary Fig. S11b**) and these cultures could be extended longer if experiment needs required.

**Figure 3.**
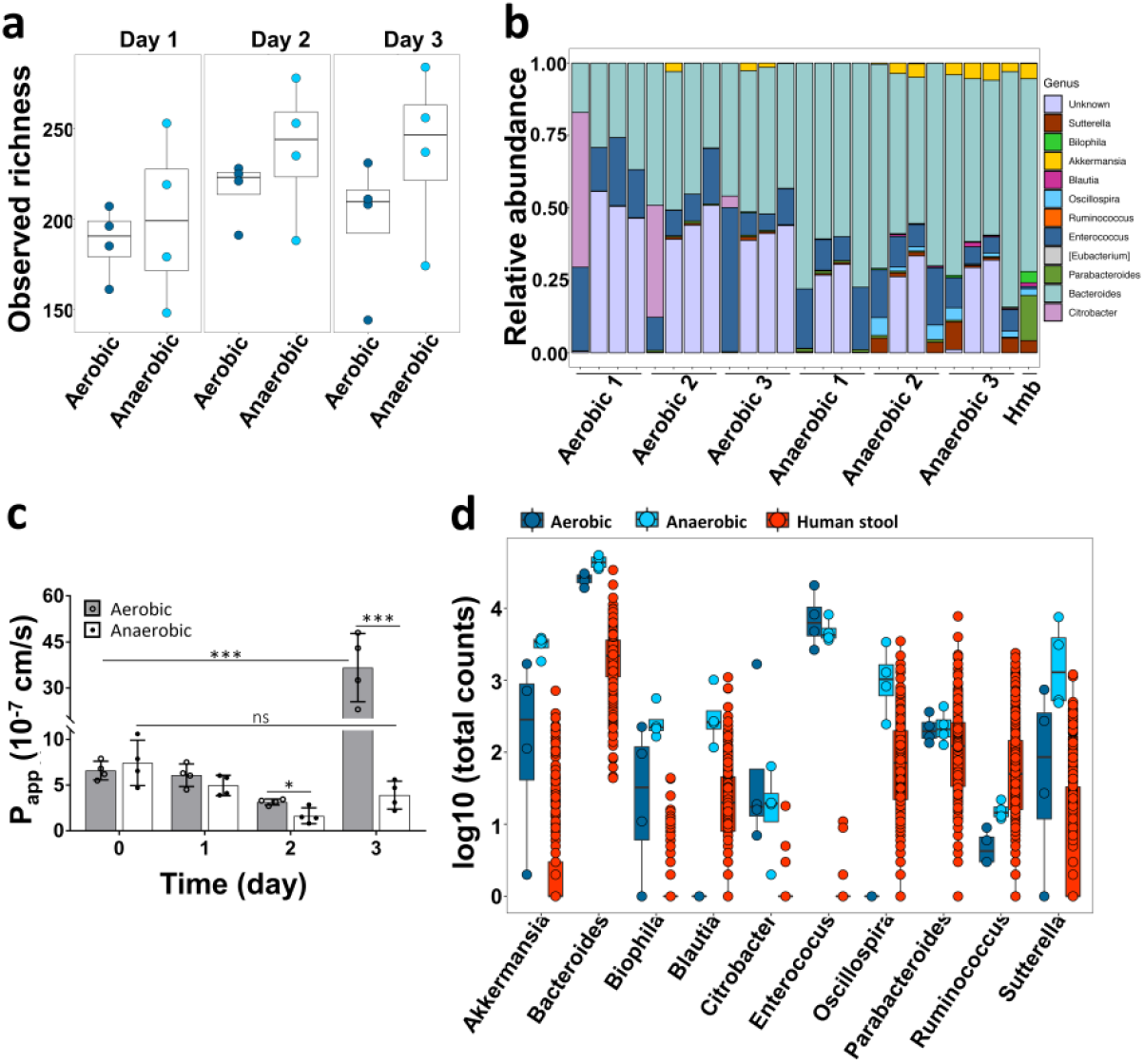
Analysis of the diversity and relative abundance of microbiota co-cultured in Intestine Chips under aerobic and aerobic conditions. (**a**) Observed alpha diversity (richness) in our complex gut microbiome samples when cultured for 1 to 3 days in direct contact with human Caco2 intestinal epithelium (each data point represents one Intestine Chip). (**b**) Relative abundance of genera measured across all samples highlighting changes in the abundance of the different genera observed over time. Data points represent each of the 3 replicates cultured under aerobic or anaerobic conditions at 0, 1, 2 or 3 days of culture (left to right, respectively); Hmb indicates genera abundance in the complex microbiome stock at time 0. (**c**) Changes in apparent paracellular permeability (P_app_) measured by quantifying cascade blue transport across the tissue-tissue interface within the Intestine Chip after co-culture with complex gut microbiome under aerobic (gray) and anaerobic (white) conditions (n=4; *, P < 0.05; ***, P < 0.001). (**d**) Differences in microbial abundance between Intestine Chip samples (dark blue: aerobic; light blue: anaerobic) and human microbiome stool sample from the Human Microbiome Project (red). Data are shown as log10 of the total number of reads; each data point corresponds to a single sample.

To further assess the physiological mimicry obtained using the anaerobic Intestine Chip lined by Caco2 epithelium, we compared the genera identified in this study with publicly available data from studies of human stool generated by the Human Microbiome Project^34^ (**Fig. 3d**). We did not expect the composition of the microbiome grown on chip to precisely recapitulate that of stool because the microbiome of the small intestine is known to show regional differences^35,36^. Nevertheless, our results show that the anaerobic culture system provides an environment for complex gut microbiota that sustains a diverse bacterial community, which falls into the range of abundances reported in the Human Microbiome Project. Furthermore, the relative abundances of the phyla that dominate the human gut, *Bacteroidetes (Bacteroidetes* and *Parabacteroides* genera) and *Firmicutes (Blautia, Enterococcus, Ruminococcus,* and *Oscillospira* genera), were higher in the anaerobic chips than in the aerobic chips with some genera (*Blautia* and *Oscillospira*) missing in the aerobic chips altogether (**Fig. 3d**). Oxygen sensor readouts in aerobic and anaerobic chips cultured with a viable microbiome or sterilely (microbe-free) confirmed that the oxygen concentration was maintained below 1% throughout 5-day co-culture period in anaerobic co-cultures (**Supplementary Fig. S11c**). Moreover, these results showed a decrease in oxygen concentration in aerobic chips cultured with microbiome over time (**Supplementary Fig. S11c**), which is similar to what we observed in the co-culture with *B. fragilis.* This was likely due to the increased vertical growth of villi we observed in these chips relative to anaerobic chips, as well as to concomitant oxygen utilization by the bacteria, which increased in numbers by day 1 in both aerobic and anaerobic chips (**Figures S11c**). Although the oxygen concentration in the aerobic chip never reached the low levels obtained in anaerobic chips, this decrease in oxygen could explain the presence of some obligate anaerobes, such as *Akkermansia*, that we observed in the aerobic chips; however, clearly the constant anaerobic conditions are more optimal for maintenance of co-cultures of anaerobic bacteria with viable human cells. Interestingly, the genus *Akkermansia,* which has been recently implicated as an enhancer of gut barrier function^37–39^, showed a considerably higher number of total counts in the anaerobic culture system compared to human stool (**Supplementary Fig. S12, S13**). Additionally, the genus *Enterococcus* was found to be present at higher levels in both chip culture systems compared to the stool samples, suggesting that some gut microbial species may grow better under conditions that more closely mimic regions of the living intestine than in stool. Taken together, these data confirm that this anaerobic human Intestine Chip system enables living human intestinal epithelium to be co-cultured in the same channel as a complex human gut microbiome containing a range of bacterial genera that come much closer to what is observed in healthy human donors than has ever been possible before.

To determine the stability of the microbial communities in the anaerobic Intestine Chip system, we analyzed their change in abundance over 3 days of co-culture with human Caco2 intestinal epithelium and underlying endothelium. Our results show that genera composed of obligate anaerobes, such as *Akkermansia, Blautia, Bilophila,* and *Suterella,* increased in abundance over time, presumably due to maintenance of low oxygen concentrations (**Fig. 4a, top**). *Bacteroides*, the highest abundance genus in the anaerobic Intestine Chips, remained relatively stable over time. Additionally, in a subsequent experiment, we cultured anaerobic Intestine Chips with the same Hmb microbiome stock for 5 days and plated serial dilutions of the flush on anaerobic Brucella culture plates in the presence or absence of vancomycin. Our results show that the bacterial load increased at day 1 compared to the seeding inoculum, and then remained constant throughout the experiment (**Supplementary Fig. S11d**). Vancomycin kills Gram positive bacteria, and thus the difference in counts between Brucella plates in the presence versus absence of vancomycin suggests both Gram positive bacteria and Gram negative bacteria (*i.e.*, that survived in the vancomycin plates) remained viable over 5 days of co-culture with the intestinal epithelium on-chip. Thus, although serial dilutions of a mixed population of bacteria do not provide accurate total counts, these data show that both Gram positive and Gram negative bacteria remained viable and proliferated (increased in number) over time in the human Intestine Chips.

**Figure 4.**
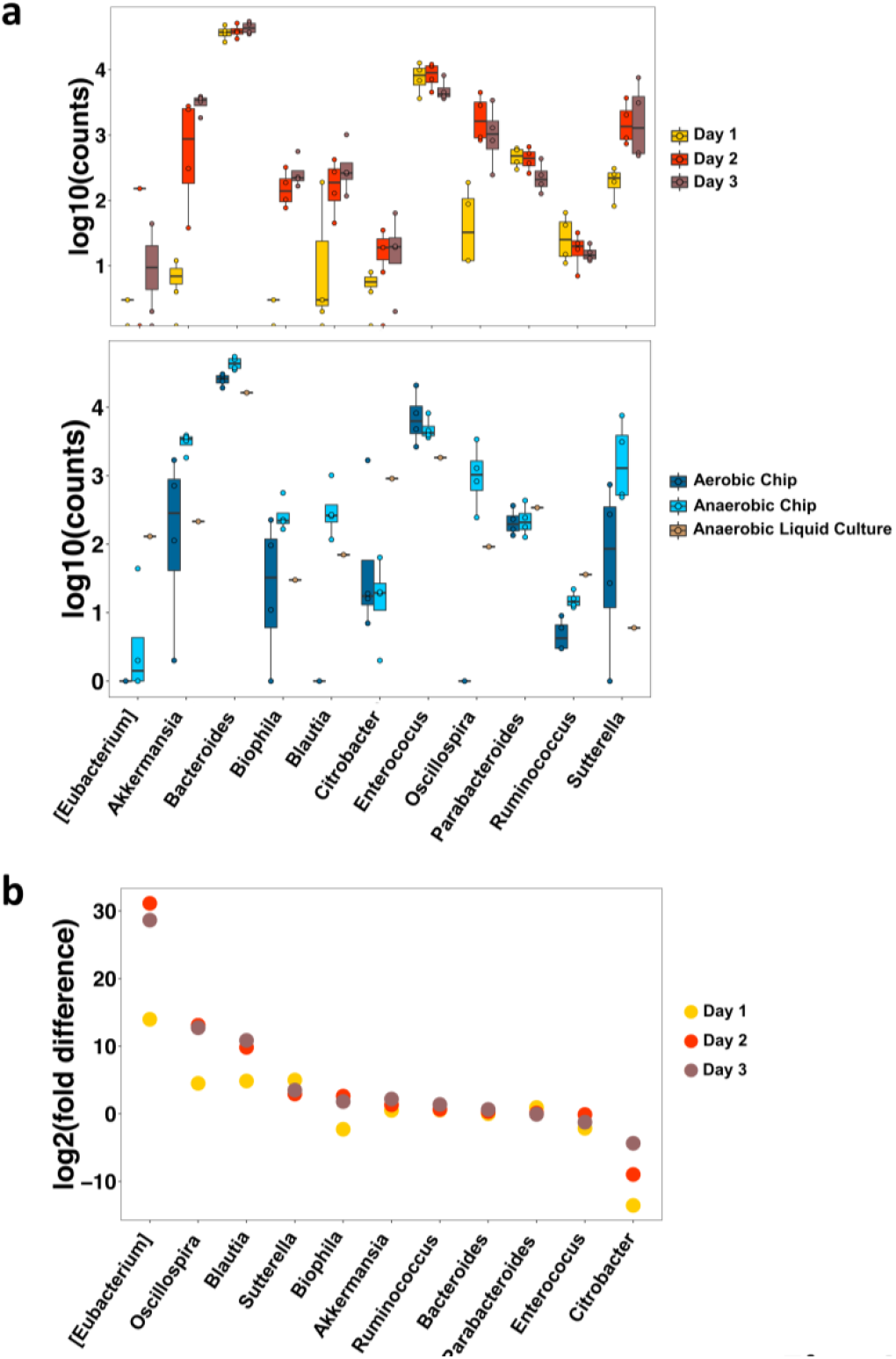
Anaerobic conditions in the Intestine Chip enhance the growth of multiple genera compared to the aerobic chip and conventional liquid culture. (**a**) Differential abundance bacterial genera in the Caco2 Intestine Chip measured under anaerobic conditions over 3 days of culture (top), or at day 3 in the anaerobic chip (light blue) compared to the aerobic chip (dark blue) or anaerobic liquid culture (bottom). Data are presented as log10 of the total read counts for each genus; each data point represents one chip. The total read counts for all genera at the bottom are normalized to their counts in liquid culture.(**b**) Differential abundance in bacterial genera measured over 1 to 3 days of co-culture in the anaerobic versus aerobic Intestine Chip. Differential abundance is represented as log2 (fold change); each data point corresponds to the differential abundance for a given genus at a given day, comparing anaerobic to aerobic cultures (n = 4 chips for each group on each day).

Importantly, when we compared the growth of microbiota cultured for 3 days in the anaerobic Intestine Chip with that produced by culturing the same microbiome samples in conventional liquid medium culture in an anaerobic chamber, we observed significantly different growth responses for multiple genera (**Fig. 4a, bottom**). Notably, the genus *Akkermansia* shows preferential growth in the anaerobic Intestine Chip, presumably due to the presence of high levels of mucus that is produced by the Caco2 intestinal epithelium on-chip^39^, which complements the mucins already present in the medium. Bacteria in the *Blautia, Bilophila, Oscillospira*, and *Suterella* genera also showed enhanced growth in the anaerobic chips containing living human intestinal epithelium compared to anaerobic liquid culture, whereas the Gram-negative obligate aerobe, *Citrobacter*, was less abundant on-chip. Thus, the presence of human intestinal epithelium that secretes a natural mucus layer above its apical surface^25^ is crucial for culturing and sustaining the complex features of human microbiome on-chip.

To complement these analyses, we calculated the differential abundance of the different genera over time in the anaerobic versus aerobic Intestine Chips. Our results show that the obligate anaerobes, *Eubacterium, Oscillospira, Blautia*, and *Suterella* were significantly more abundant in the anaerobic chips compared to aerobic chips over our time course (FDR q < 0.05), whereas the obligate aerobe, *Citrobacter,* consistently showed a lower abundance in the anaerobic chip (**Fig. 4b**). Whether taking into account the abundance of the various genera in anaerobic liquid culture (**Supplementary Fig. S12**) or in the Hmb microbiome stock (**Supplementary Fig. S13**), quantification of the total read counts confirmed that the total numbers of obligate anaerobes, including *Sutterella, Blautia, Oscillospira, Bilophila*, and *Akkermansia*, were significantly higher in the anaerobic chips. Taken together, these results confirm that the hypoxia gradient system combined with the presence of a living human intestinal epithelium provides a unique and preferential environment for sustained culture of anaerobic as well as aerobic gut bacteria from diverse genera.

Finally, we explored whether this experimental approach can be used to co-culture complex gut microbiome obtained from *fresh* human stool specimens in direct contact with *primary* human intestinal epithelium (*i.e.*, rather than using the established Caco2 intestinal cell line). To do this, we engineered human Intestine Chips lined with intestinal epithelial cells isolated from organoids derived from normal regions of surgical biopsies of human ileum, which exhibit multi-lineage differentiation, villi formation, and mucus production when grown on-chip^27^. We then inoculated the epithelial channels of 4 different chips with complex microbiome isolated from fresh human stool samples collected from four different infants (one with a corrected gestational age of 30 week and three with an age of 36 week). DIC (**Fig. 5a**) and confocal fluorescence microscopic (**Fig. 5b**) imaging of the primary human Ileum Chips confirmed the presence of a villus intestinal epithelium lined by a continuous polarized epithelium with an F-actin-containing and villin-stained brush borders along its apical membrane and basal nuclei. As expected, the bacterial richness was reduced in the infant stool stock (586 OTUs) compared to adult human-derived stool (938 OTUs) at the same dilution per gram of stool, and these differences in richness were accurately recapitulated on-chip. We found that the primary human intestinal epithelium could be co-cultured in direct contact with this complex gut microbiome without compromising epithelial barrier function, and this co-culture was stably maintained for up to 5 days on-chip (**Fig. 5c**), much as we had observed with the Caco2 epithelium. Importantly, the microbiome cultured in these primary Intestine Chips also maintained a high bacterial richness, ranging from 118 to 135 OTUs (**Fig. 5d**) corresponding to 6 phyla (*Actinobacteria, Bacteroidetes, Cyanobacteria, Firmicutes, Proteobacteria and Tenericutes*) and 32 unique genera. Thus, the Intestine Chip method can be used to sustain a complex community of human microbes in direct contact with normal, patient-derived, human intestinal epithelial cells for many days in culture.

**Figure 5.**
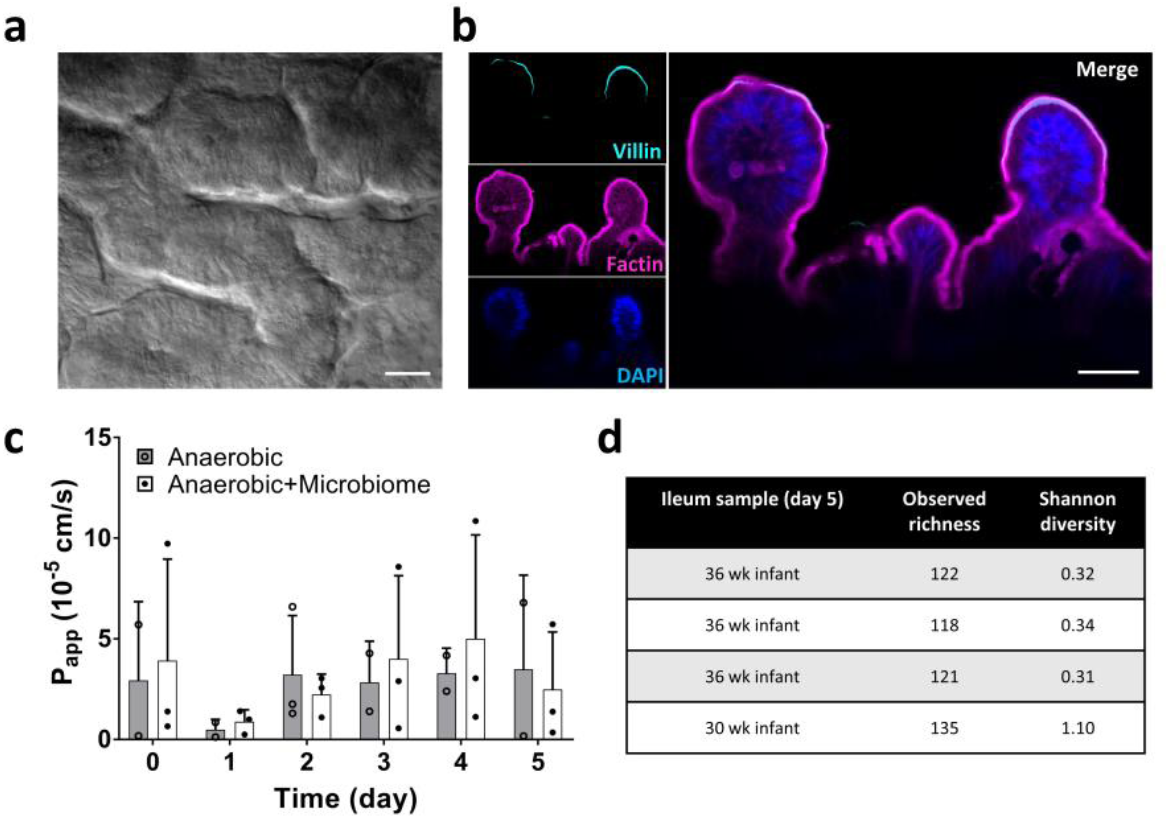
Anaerobic co-culture of gut microbiome obtained from fresh human patient-derived stool with primary human ileal epithelium in the Intestine Chip. (**a**) Microscopic views showing the villus morphology of the primary ileal epithelium cultured for 5 days in the Intestine Chip under anaerobic conditions when viewed from above by DIC (bar, 50μm) or (**b**) shown in cross-section by confocal immunofluorescence imaging for villin (cyan, top), F-actin (magenta, middle) and DAPI (blue, bottom; bar, 50μm). (**c**) Changes in apparent paracellular permeability (P_app_) measured by quantifying cascade blue transport across the tissue interface within the primary Intestine Chip during co-culture with or without complex human gut microbiome under anaerobic conditions (bacteria contained with patient-derived stool samples were added on day 0). (**d**) Observed alpha diversity (richness) and Shannon diversity of bacteria measured in effluent samples from the epithelial channel of primary Intestine Chips after 5 days of co-culture with stool microbiome samples collected from 30 or 36 week old neonates (4 different individuals).

## DISCUSSION

Given the importance of commensal gut microbiome for human health and the lack of any *in vitro* model that can faithfully mimic the complex epithelial-microbe interactions that occur across the host-microbiome interface, we leveraged human Organ Chip technology to develop a device that enables human intestinal epithelium to be co-cultured with the highly diverse community of commensal microbes that comprises the human gut microbiome under aerobic and anaerobic conditions. Our results show that the anaerobic human Intestine

Chip offers a robust modular platform for recapitulating the human intestinal-microbiome interface *in vitro*. Using this device, for the first time, it is possible to stably co-culture a complex living microbiome in direct contact with living mammalian cells for 5 days or more *in vitro*. This is significantly longer than past studies using the HMI model that only sustained co-cultures of intestinal epithelium with complex microbiome for 48 hours, and in which the bacteria had to be physically restricted from contacting the epithelium by a semi-permeable membrane to ensure epithelial viability^17,40^. Importantly, providing a physiologically-relevant low oxygen microenvironment on-chip also sustained a higher level of microbial diversity (∼200 unique OTUs), increased abundance of obligate anaerobic microbiota compared to aerobically-cultured chips, and maintained a diverse community of commensal microbes that closely resembled that of the human gut microbiome *in vivo*. For example, when the complex gut microbiome was cultured in the anaerobic Intestine Chip, it maintained an abundance of obligate anaerobic bacteria with ratios of *Firmicutes* and *Bacteroidetes* similar to those observed in human feces^41^.

Using a custom-designed anaerobic chamber and chips containing oxygen sensors that enable monitoring of local oxygen concentrations on-chip, we were able to recapitulate *in vivo*-like oxygen gradients, which also allowed us to demonstrate morphological and functional changes in the intestinal epithelium in response to altered oxygen levels. When the epithelium was co-cultured on-chip with either the obligate anaerobe *B. fragilis* or complex human microbiome under anaerobic conditions, we observed bacterial growth that was accompanied by enhanced intestinal barrier function compared to aerobic chips. Moreover, a similar enhancement of intestinal barrier function was obtained when we cultured complex gut microbiome under anaerobic conditions, which again is consistent with *in vivo* findings^39^.

Oxygen tension is one of the main regulators of intestinal function and pathogenesis of GI diseases^42,43^. By integrating non-toxic oxygen sensors into our devices, we were able to measure oxygen levels throughout the microfluidic Intestine Chips without interference with microscopic imaging, device fabrication or cell culture. Use of these sensors, rather than incorporating multiple external oxygen-detecting probes, enables this approach to be more easily scaled to create many miniaturized Organ Chip platforms. The anaerobic chamber we engineered also generates radial oxygen gradients across the endothelium-epithelium-microbiome interface that allows oxygenation of the human tissues while providing an anaerobic environment for growth of the obligate anaerobes. Anaerobic incubators or glove boxes can be used to maintain hypoxic conditions for bacterial cultures, but they commonly provide a single uniform low oxygen concentration, rather than physiologically-relevant oxygen gradients directed across tissue-tissue interfaces. In contrast, our anaerobic chamber is portable, highly customizable, compatible with imaging, and most importantly, capable of engineering oxygen gradients across the endothelial-epithelial interface of any Organ Chip on demand.

Oxygen concentrations in the lumen of the human intestine are known to affect the spatial distribution and metabolism of gut flora^44^, and most intestinal bacteria are obligate anaerobes, many of which fail to grow at oxygen concentrations greater than ∼0.5%^45^. Any culture system that is designed to recapitulate the host gut-microbiome interface must therefore be able to achieve and sustain oxygen concentrations at these low levels. A microfluidic-based anaerobic culture system has been described previously that maintains oxygen levels as low as 0.8% in the presence of a facultative anaerobe^40^, but this level is still too high to support obligate anaerobes. Moreover, this model used both a synthetic mucus layer and a nanoporous membrane to physically separate bacteria from the intestinal epithelium^40^. Using our custom anaerobic chamber, we were able to attain an oxygen concentration of less than 0.3% in the epithelial channel where we cultured the commensal microbes, which is much closer to that found in the gut lumen *in vivo*^46^. Additionally, we validated the relevance of these hypoxic culture conditions by showing that they support the growth of the obligate anaerobe *B. fragilis* that cannot grow in the presence of greater than ∼0.5% dissolved oxygen^31,47^, whereas most of these bacteria died off after 3 days of *in vitro* culture under conventional aerobic conditions. Furthermore, the finding that co-culture of the human intestinal epithelium with *B. fragilis* under anaerobic conditions also increased (rather than decreased) intestinal barrier function on-chip is consistent with the finding that oral delivery of *B. fragilis* corrects intestinal permeability defects in a mouse autism model^48^.

More importantly, we found that the hypoxic human Intestine Chip model supports co-culture of complex human microbiota composed of over 200 unique OTUs and at least 11 different genera of bacteria for at least 3 days in co-culture. Bacterial members of the *Bacteroidetes* and *Firmicutes* phyla, and to a lesser degree *Verrucomicrobia* and *Proteobacteria*, which comprise the human intestinal microbiome *in vivo*^49^, also colonized our Caco2 Intestine Chips at similar ratios. Anaerobic chips had increased levels of anaerobic *Clostridia, Bacteroides*, and *Akkermansia*, whereas *Proteobacteria,* which accumulate mainly at more oxygenated regions of the proximal gastrointestinal tract^46,50^ dominated the aerobic chips. One limitation of our approach is the need to dilute the complex microbiome inoculum to avoid rapid unrestrained bacterial overgrowth. This may result in exclusion of some rare bacteria; however, this could be ameliorated by using larger Intestine Chips, optimizing the lumen perfusion rate, applying cyclic (peristalsis-like) mechanical deformations that suppresses growth of commensals^21^, or altering medium conditions to limit bacterial overgrowth. Nevertheless, these data show that the anaerobic system promoted more bacterial diversity than the aerobic system. Moreover, the anaerobic human Intestine Chip supported a wide range of bacterial genera similar to those found in human stool, which is much more complex than any microbiome community that has been previously cultured in direct contact with mammalian cells, and we could sustain these co-cultures for at least 3 to 8 days *in vitro*.

Others have previously maintained complex microbiota in test tube cultures^51^, however, our results indicate that the presence of a more *in vivo*-like intestinal tissue microenvironment significantly influences the composition of the microbial community. For example, the mucus degrading, obligate anaerobe genus *Akkermansia* was found in higher abundance in the anaerobic Intestine Chips which contain human intestinal epithelial cells that secrete mucus than in similarly anaerobic liquid cultures that were artificially supplemented with mucin. In contrast to liquid cultures, the anaerobic Intestine Chip also allows inferences to be made regarding the effects of commensal microbes on the host epithelium and vice versa. It is interesting that the enhanced growth of *Akkermansia* in the anaerobic Intestine Chip was accompanied by increased intestinal barrier function since the high abundance of this organism has been previously suggested to enhance gut barrier function *in vivo*^37–39^. While *Akkermansia* was also found in the aerobic chips, it displayed significantly lower read counts suggesting that while it increases in numbers over time in the anaerobic chips, it either cannot grow or it is no longer viable in the aerobic chips.

HIF-1α is believed to control barrier integrity by regulating multiple barrier-protective genes, and its dysregulation may be involved in GI disorders^52,53^. Interestingly, although we observed elevated HIF-1α expression in anaerobic Intestine Chip, we did not detect any changes in barrier function unless we also co-cultured complex microbiota. This system allows us to parse out which physiological effects are due to changes in oxygen levels alone and which are due to the presence of specific microbes. Defining the causative relationship between the abundance of each individual genus of bacteria and distinct functions of the co-cultured human intestinal epithelium is beyond the scope of this study. However, this system could be harnessed to address these types of questions, as well as how different commensal microbes contribute to the pathophysiology of various gastrointestinal diseases^54^ in the future.

The purpose of this study was to develop an anaerobic method for co-culturing human epithelial cells with complex human microbiome in an organ-relevant microenvironment *in vitro*. We demonstrated this capability using both an established human Caco2 intestinal cell line and primary human ileal intestinal epithelium, however, the same methodology could be applied to study host-microbiota interactions in any Organ Chip (*e.g.*, lung, skin, etc.). Furthermore, by integrating primary epithelial cells from intestinal biopsies as we did here, or patient-derived induced pluripotent stem (iPS) cells^55^, in combination with microbiomes obtained from the same patients, it should be possible to develop patient-, disease-, and location-specific, host-microbiome co-culture models. The modular nature of the Organ Chip technology also allows for the incorporation of additional cell types. In this study, we incorporated intestinal endothelium in our Intestine Chips because it enhances intestinal barrier function, villi development and mucus production^26,27,56^, but other cell types, such as immune cells and pathogens that play crucial roles in host gut-microbiome interactions^57,58^ could be incorporated as well. Our oxygen sensing chips also have the potential to be combined with on-chip TEER technology^59^ for real-time monitoring of intestinal barrier function in the presence of different cell types and individual strains of bacteria. Thus, this methodology could be used in the future to unravel complex functional links between intestinal epithelial cells, immune cells, and gut microbes to understand mechanisms of human disease, discover new therapeutics, and advance personalized medicine.

## Acknowledgements

This research was supported by the U.S. FDA grant (HHSF223201310079C), DARPA THoR grant (W911NF-16-C-0050), Bill & Melinda Gates Foundation, Wyss Institute for Biologically Inspired Engineering at Harvard University, and Fundação para a Ciência e a Tecnologia (FCT) Portugal (project PD/BD/105774/2014 to the Institute for Bioengineering and Biosciences). We thank D. E. Achatz (PreSens Precision Sensing GmbH, Germany) for graciously providing oxygen sensing particles and her expert technical advice, and T. Ferrante for his assistance with imaging.

## Author contributions

S.J-F., E.L.C., F.S.G, J.M.S.C., R.N. and D.E.I. designed the research. S.J-F, E.L.C., F.S.G., B.N., C.F., A.T., and M.C. performed experiments. S.J-F., D.M.C, E.L.C., F.S.G, B.N., D.L.K., R.N. and D.E.I. analyzed and interpreted the data. K.E.G helped in preparation of infant microbiota. D.T.B established and prepared human ileal organoids. S.J-F, F.S.G, E.L.C, D.M.C., and D.E.I wrote the paper with input from B.N., O.L., J.M.S.C., and R.N.. The paper has been reviewed, discussed and edited by all authors.

## Author information

The authors declare competing financial interests: D.E.I. holds equity in Emulate, Inc., consults to the company, and chairs its scientific advisory board. Correspondence and requests for materials should be addressed to D.E.I.

